# Saline-free preparation for chronic *in vivo* imaging in adult *Drosophila*

**DOI:** 10.64898/2026.02.18.706199

**Authors:** Ruibao Zhu, Salah Khorbtli, Jingmin Zhang, Zhuchen Fu, Cheng Huang

**Author notes:** Technical contact (Ruibao Zhu).

## Abstract

Longitudinal brain imaging is essential for understanding neural mechanisms. Here, we present a saline-free, chronic preparation for repeated neural recording in adult *Drosophila* over multiple days. We describe steps for mounting flies, performing manual surgery on the head cuticle without external saline, and resealing the opening to create a transparent optical window. We demonstrate the utility of this approach by tracking single-neuron spiking and neuronal calcium dynamics over 7–10 days. This protocol is potentially applicable to other insect species.

**Graphical abstract:** 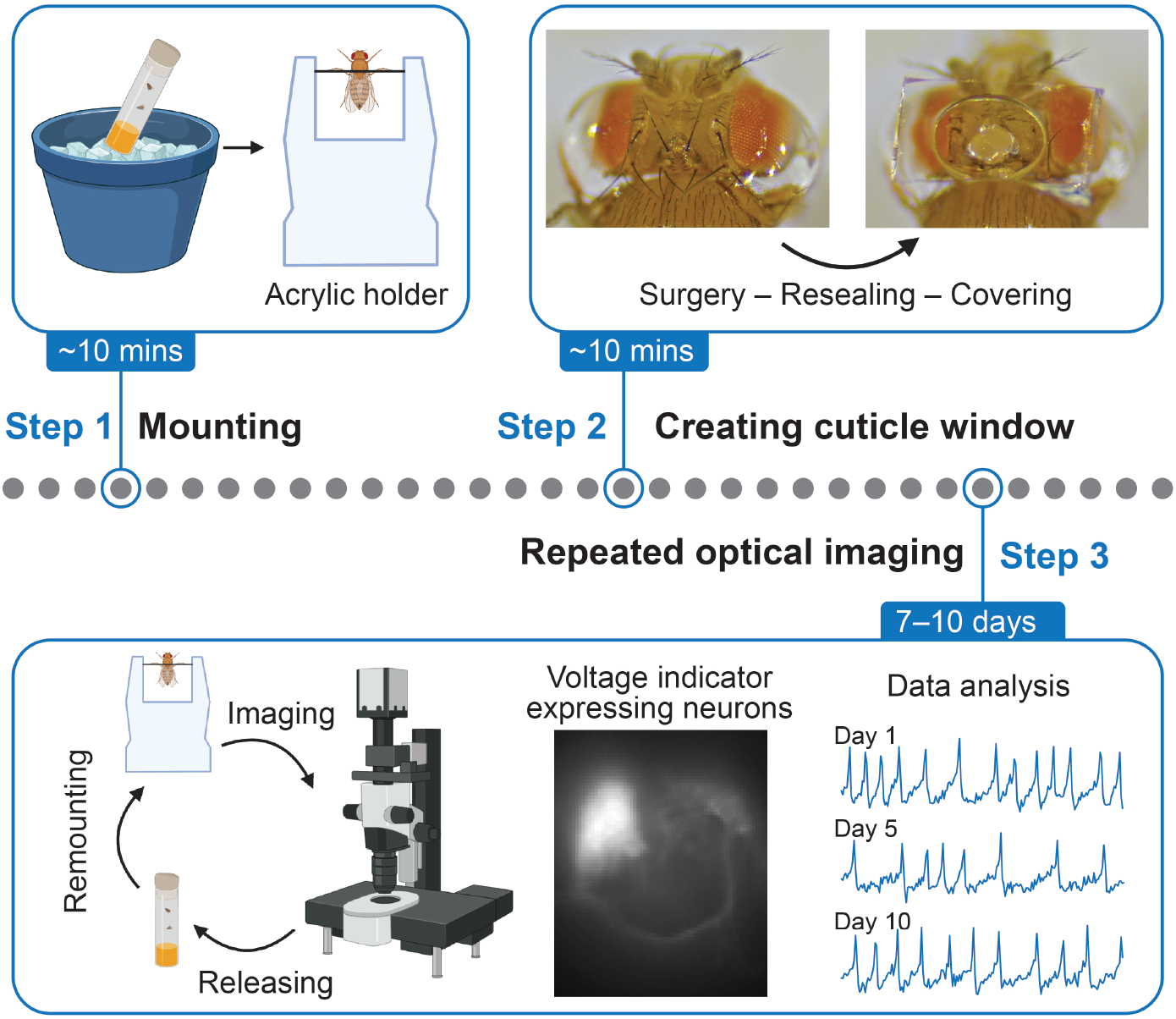

## Before you begin

### Background

Understanding the mechanisms of neural function, plasticity, development, and age-related changes requires direct, time-lapse observation of cellular and physiological events within living organisms over extended periods. In many animal models, chronic preparations for long-term intravital microscopy have revealed rich longitudinal dynamics that are often missed in studies of different individuals at discrete time points. *Drosophila melanogaster* is a cornerstone for biological research due to its sophisticated genetic toolkit, but achieving stable, long-term *in vivo* imaging of its central nervous system, especially for recording neural activity over multiple days, has remained a significant challenge. The fly’s inherent fragility and opaque exoskeleton limit most physiological recordings to brief durations of a few hours.

Removal of the head cuticle is the essential step for optical access to the *Drosophila* brain. The widely adopted dissection approach for *in vivo* physiology requires submerging the exposed brain in an aqueous saline solution to maintain osmotic and ionic balance^1,2^. While effective for acute or short-term experiments, this saline-based approach presents significant limitations for extended imaging sessions. Typically, recording durations range from 1 to 24 hours^3,4^. While recent laser-based microsurgery techniques can offer high precision for cuticle removal without the need for external saline and enable reliable recordings for several weeks^5,6^, the considerable technical demands, including specialized and costly equipment, limit their widespread adoption and accessibility.

To overcome these limitations, we simplified the previous laser microsurgery approach^5^ to develop a manual, saline-free, chronic preparation for adult *Drosophila*. This optimized manual surgical approach enables repeated recording of neural activity over 10 days. This protocol describes a complete workflow, including fly mounting, manual dissection, and cuticle resealing steps to create a durable, transparent optical window. Crucially, these procedures are performed without the need for external saline during surgery or subsequent imaging sessions. We demonstrate the utility of this approach by tracking single-neuron spiking activity and neuronal calcium dynamics in the adult fly brain over periods of 7–10 days.

### Preparation of the tethering station and fly holder

Before starting the mounting process, prepare a custom-built tethering station (Figure 1A), an acrylic fly holder (Figure 1B) installed on a metal bar (Figure 1C), UV-curable glue (Figure 1D), and surgical tools (Figure 1E). The design of the tethering stage is based on previous work by the Reiser group^7^ with modifications. Detailed assembly instructions and a parts list are available in the online tutorial: https://flymemorylab-washu.github.io/Tethering-Station-Assembly.

**Figure 1.**
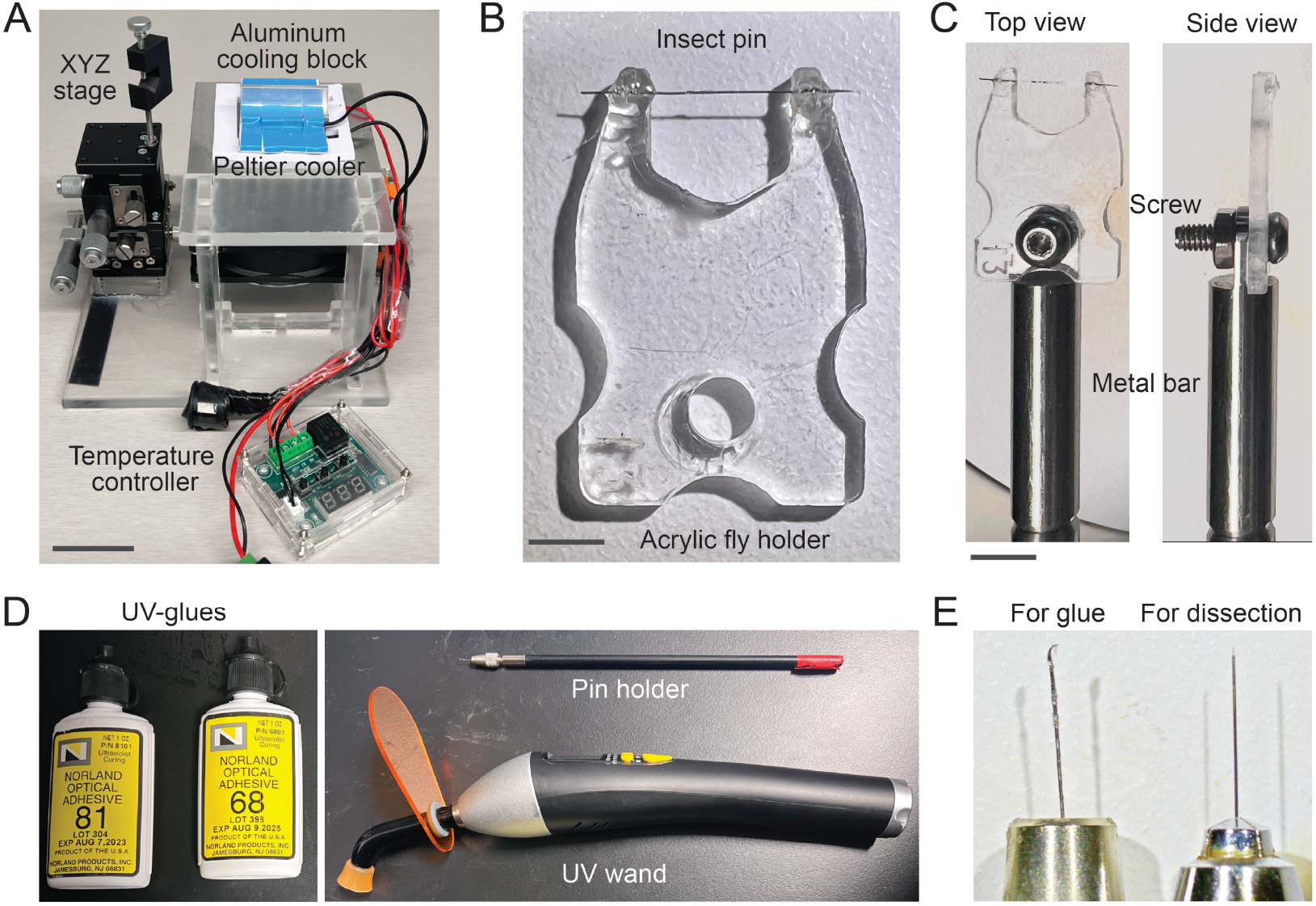
Fly tethering station and surgical tools. (A) Assembled tethering station with a multi-axis stage and a cooling block. Scale bars: 4 cm. (B) Example of a custom-built acrylic fly holder. Scale bars: 4 mm. (C) Top (*left*) and side (*right*) views of a fly holder installed on a metal bar. Scale bars: 6 mm. (D) Examples of the UV-curable epoxy and UV wand needed. (E) Surgical needles used for applying glue (Minutien pins with nickel-plated pin holder; *left*) and for dissecting cuticle (Tungsten needle with Moria pin holder; *right*).

### Drosophila husbandry

To establish a *Drosophila* stock for imaging, cross a selected GAL4 line (or another expression system) with the desired fluorescent reporters (e.g., voltage, calcium, or other indicators). Raise the flies on standard cornmeal agar food under a 12-hr light/dark cycle at 25°C and 50% relative humidity. Select 5-to 8-day-old flies from the vials at the time of surgery.

### Innovation

This saline-free approach minimizes tissue damage, maintains physiological consistency, and enhances animal viability, enabling reliable, chronic in vivo observations. Compared with prior complex laser-based approaches^5,6^, this simplified manual preparation is cost-effective and suitable for wider adoption across the *Drosophila* research community. Furthermore, this protocol may also be extended to long-term brain imaging in other insect species with minimal modifications.

### Key resources table

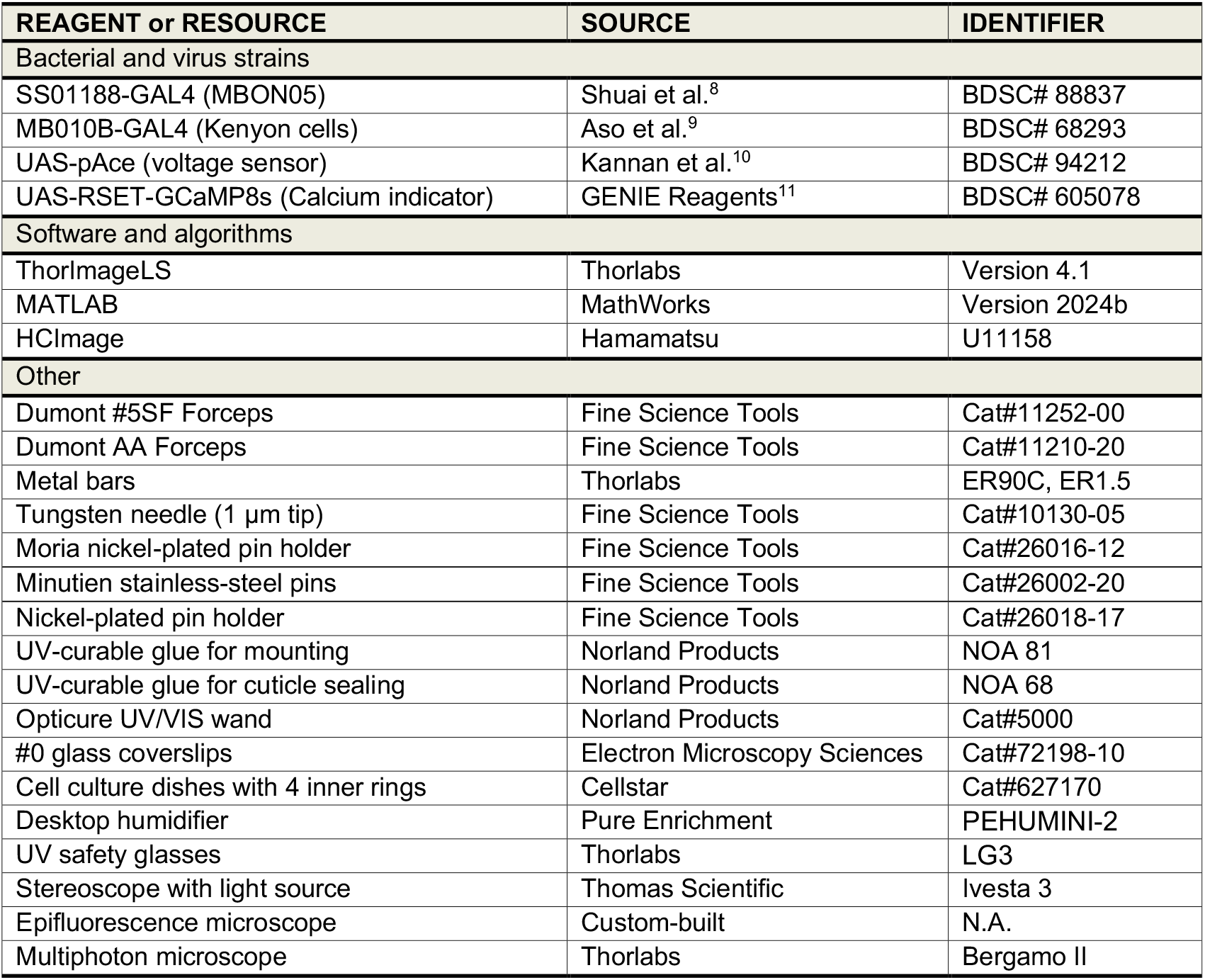

## Step-by-step method details

### Mounting flies to the acrylic holders

#### Timing: 5–10 min per fly

This section outlines the steps for mounting a fly to a rigid holder to minimize motion artifacts during surgery or imaging while ensuring the animal remains awake and healthy.

1. Install an acrylic fly holder on the metal bar using a screw (Figure 1C), then clamp the bar to the multi-axis stage of the tethering station (Figure 1A).
2. Anesthetize selected flies by placing the vial on ice for 1 min.
3. Transfer an anesthetized fly onto the cooled aluminum surface (∼4°C) of the thermoelectric cooling block (Figure 2A).
4. Using dull forceps (Dumont AA), gently adjust the fly’s orientation under the stereoscope (Ivesta 3) so that the fly’s ventral side faces the aluminum block (Figure 2B). **NOTE:** Position the fly’s proboscis against a flat washer (Size M2, 5 mm outer diameter) on the aluminum block. This helps maintain the fly’s natural standing posture.
5. Apply approximately **1 µL** of UV-curable epoxy (NOA 81; Figure 1D *left*) to the middle point of the pin on the acrylic fly holder. Use the multi-axis stage to move the acrylic fly holder directly above the fly’s thorax (Figure 2C). **NOTE:** Use the insect pin with a bent tip to dip and transfer the epoxy (Figure 1E *left*). **NOTE:** Alternatively, the glue can be applied to the pin before positioning the fly.
6. Lower the holder and bring the pin into contact with the dorsal side of the fly’s thorax. Let the epoxy spread over the posterior thorax cuticle.
7. Cure the epoxy with the UV wand (Norland Inc.) for **5 s** to create a stable bond (Figure 2D). **NOTE:** UV exposure time and distance must be carefully calibrated. In our setup, we use 365/385 nm wavelengths for **5 s** with the UV wand tip approximately **2 cm** away from the fly. **NOTE:** Adjust the distance between the UV wand and the fly to prevent overheating (which can be lethal) or incomplete solidification of the glue.
8. To stabilize the head, apply approximately **0.5 µL** of UV epoxy onto each side of the fly’s head and secure the head to the thorax (Figure 2E). Cure the glue with UV light. **NOTE:** For optimal access to the target neuron population, the head orientation should be fine-tuned with the micromanipulator. Moving the fly’s body toward or away from the washer changes the head pitch angle. For imaging anterior brain regions, fix the head as shown in Figure 2D; for imaging posterior structures such as Kenyon cells, fix the head as shown in Figure 2B.
9. After mounting, allow flies to recover for **1 min** before the surgery. Ensure the head-fixed fly is fully awake and active on the holder **NOTE:** See Methods video S1 for the full mounting procedure.

**Figure 2.**
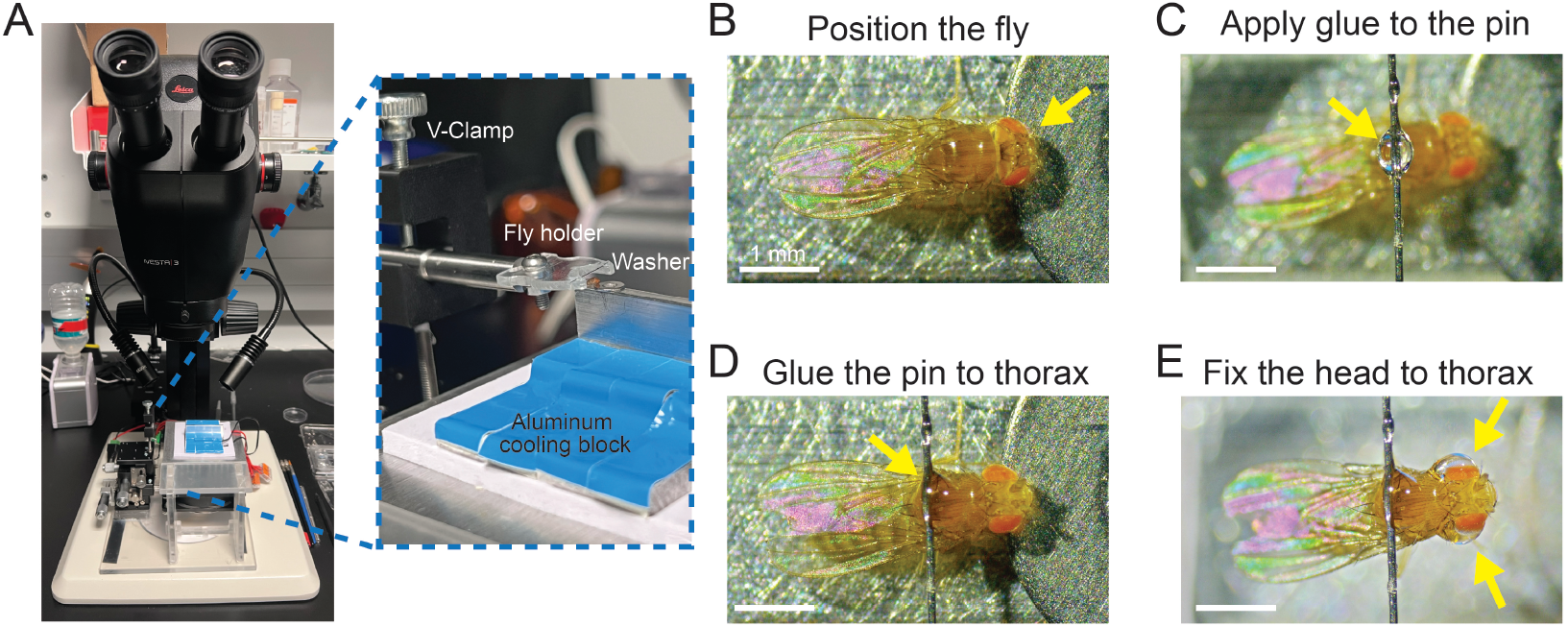
Fly mounting procedure. (A) The tethering station is placed under a stereoscope for fly mounting. The inset shows a zoomed-in view of the installed fly holder and the cooling block. (B) Top view of the fly positioned on the cooling surface with its head supported by the washer (yellow arrow). (C) A drop of UV glue applied to the pin (yellow arrow). (D) Top view of the fly with its thorax fixed to the pin (yellow arrow). (E) The fly’s head is glued to its thorax from both sides (yellow arrows). Scale bars: 1 mm in B–E.

## Saline-free surgical process

### Timing: 5–10 min per fly

This section details the surgical procedures to expose the fly brain and create a transparent window for imaging.

10. Move the tethering station away and place the acrylic holder with the mounted fly under the stereoscope (Figure 3A).
11. Use sharp Dumont #5SF forceps to carefully shorten or remove all bristles on the cuticle over the target area on the fly’s head (Figure 3B). **NOTE:** Removal or shortening of bristles is essential to prevent obstruction during imaging.
12. Use the tungsten needle (Figure 1E *right*) to gently poke multiple small holes along the perimeter of the intended optical window on the cuticle. **NOTE:** Do not penetrate too deeply to avoid damaging the underlying brain tissue. **NOTE:** A typical window size is ∼ 0.3–0.4 mm in diameter. **NOTE:** If ambient humidity is below 35%, use the desktop humidifier to maintain the fly brain’s hydration during surgery.
13. Using sharp forceps, carefully peel off the perforated cuticle to expose the brain (Figure 3C).
14. Gently clean any remaining air sacs and fat tissue that float in the hemolymph around the exposed area (Figure 3D). **NOTE:** Do not scrape the brain surface; only skim the floating tissue. Some transparent tissues may be observed in deeper areas, but do not remove them, as they will not affect imaging.
15. Apply approximately **1 µL** of UV-curable epoxy (NOA 68) to cover the entire opening on the fly’s cuticle (Figure 3E), and cure the epoxy with UV light for **5 s**. **NOTE:** A distinct light reflection around the edge of the sealed opening indicates successful sealing.
16. For experiments requiring a water-immersion objective, glue a small piece of #0 coverslip (cut into ∼1 mm × 0.5 mm pieces) onto the sealed optical window by adding a secondary layer of UV epoxy (NOA 68) (Figure 3F).
17. If necessary, apply additional UV epoxy to raise the glue height and create adequate space between the fly’s antennae and the coverslip.
18. Cure the additional epoxy with UV light. **NOTE:** See Methods video S2 for the surgical procedure.

**Figure 3.**
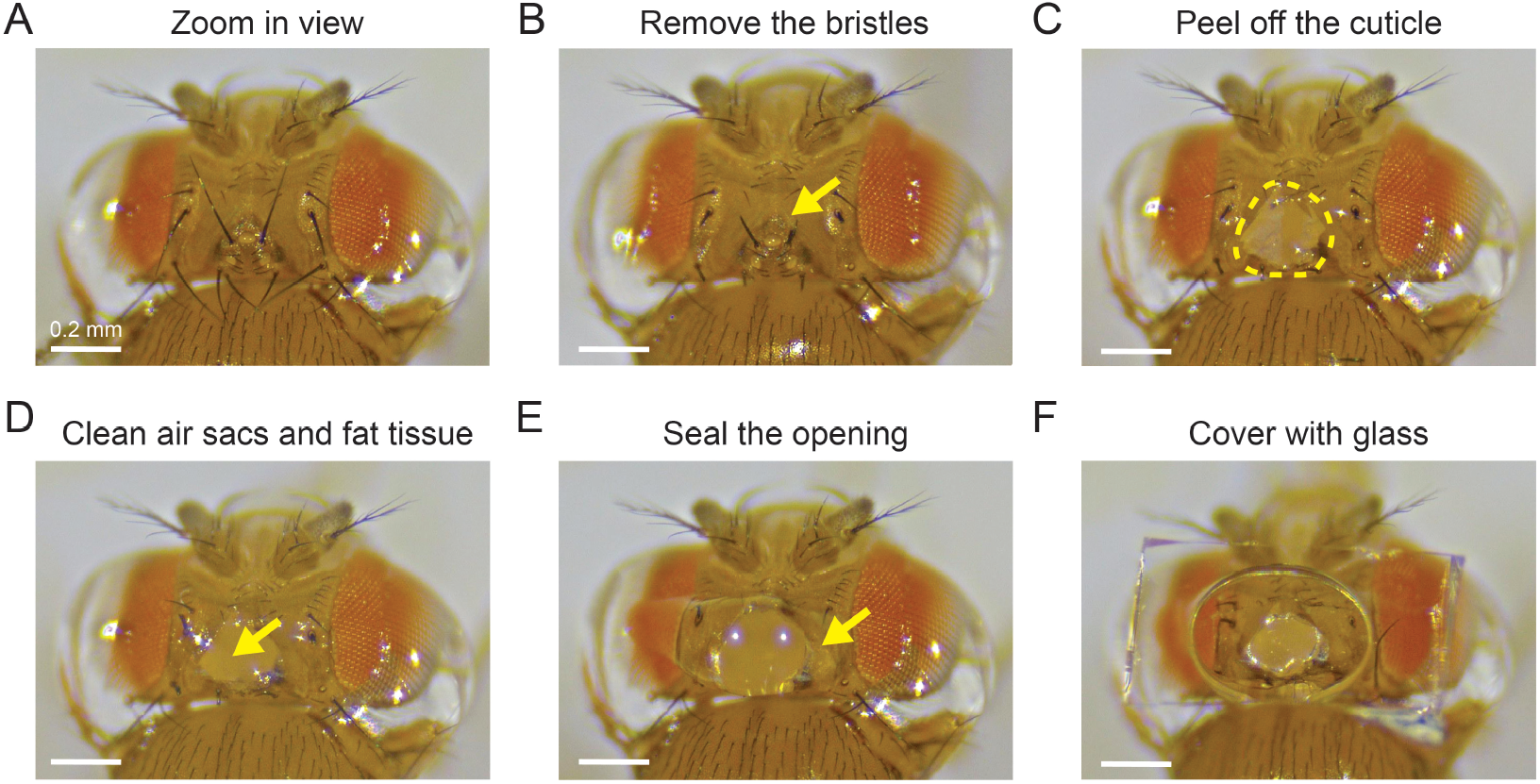
Surgical procedure. (A) Example view from the stereoscope of the tethered fly. (B) Bristles on the cuticle were removed or shortened. (C) After peeling off a piece of cuticle, the underlying tissues were exposed (*yellow outline*). (D) After removing the air sacs and fat tissues (*yellow arrow*). (E) The opening was resealed with the UV-glue (*yellow arrow*). (F) A small piece of coverslip is installed on top of the optical window. All scale bars: 0.2 mm.

## Longitudinal fluorescence brain imaging

### Timing: 1–10 days

This section describes the main steps for longitudinal imaging through the transparent window.

19. Place the acrylic fly holder onto the sample stage and position it under the fluorescence microscope (Figure 4A). **NOTE:** This procedure is compatible with most upright microscopes. For demonstration purposes, we utilized a custom-built epi-fluorescence microscope for voltage imaging and a Thorlabs Bergamo II two-photon microscope for calcium imaging. **NOTE:** When using the fly-on-a-ball system for neural activity imaging, first install the acrylic fly holder onto the mounting bar, then clamp the bar onto the multi-axis stage.
20. Apply approximately **2 µL** of distilled water to the tip of the water-immersion objective (Figure 4A).
21. Carefully lower the objective until the water droplet contacts the coverslip above the fly’s head (Figure 4B and 4C).
22. Perform fluorescence imaging using your imaging station.
23. For longitudinal imaging, ensure consistent region-of-interest (ROI) selection across different days.

**Figure 4.**
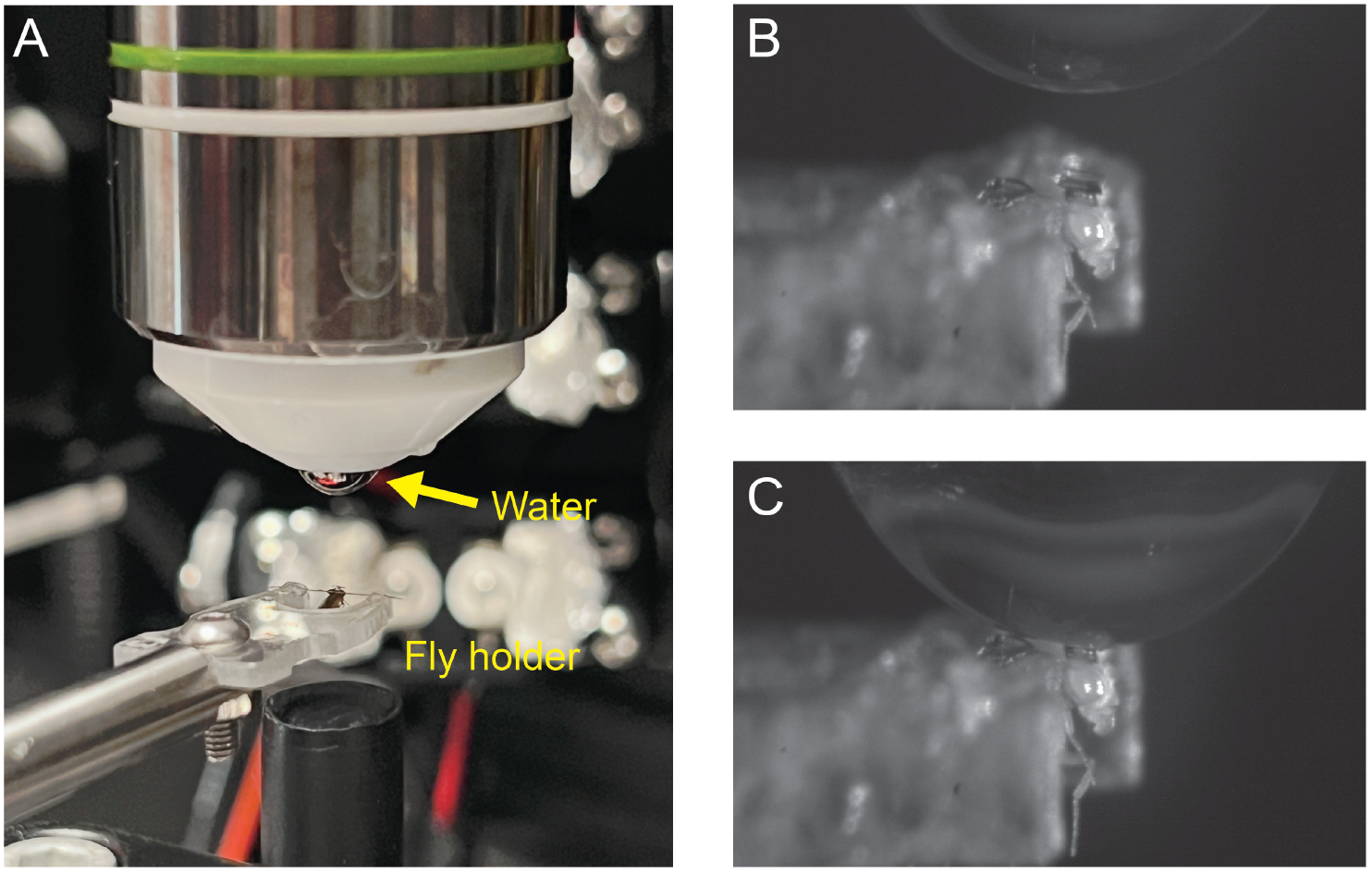
Imaging procedure. (A) The fly holder was placed under a water-dipping objective with water applied on its tip (*yellow arrow*). (B) Close view of the water droplet on the objective and the fly. (C) Lower the objective to make droplet contact with the glass.

## Fly husbandry after imaging sessions

### Timing: 5 min per fly

This section describes the steps to release the fly back to food after each imaging session.

24. After each imaging session, place the mounted fly under the stereoscope to release the fly. Using the tip of a pair of dull forceps, gently push the coverslip to break the piece of cured UV epoxy that secures the fly to the pin on the fly’s thorax (Figures 5A and 5B; See Methods video S3).
25. Once released, use the forceps to gently grasp the fly by its wings and transfer it back to a food vial (Figure 5C).
26. For subsequent imaging sessions, remount the fly by repeating the initial mounting procedure. **NOTE:** Remounting reduces restraint stress but adds repeated cold anesthesia. **NOTE:** Alternatively, to avoid repeated anesthesia and remounting, the fly can remain on the holder for extended periods. To maintain viability, feed the fly three times per day by applying 100 mM sucrose water directly to its proboscis using a micropipette or Kimwipes tissue.

**Figure 5.**
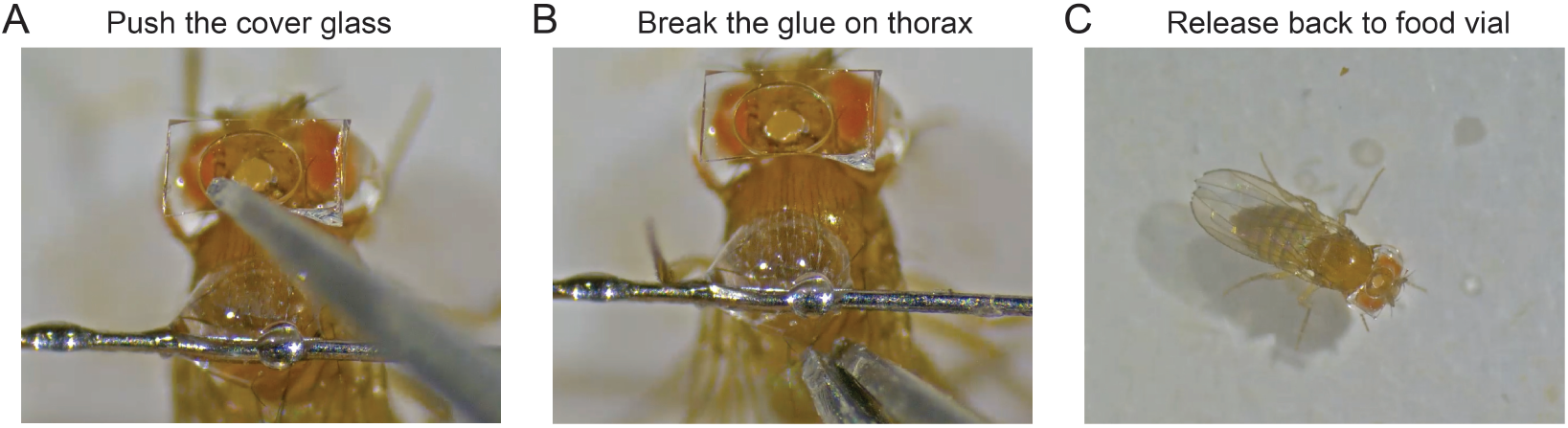
Releasing procedure. (A) Pushing the glass with dull forceps. (B) Break the glue on the thorax. (C) The fly is released back to a food vial.

## Expected outcomes

### Longitudinal voltage imaging with single-spike resolution across 10 days

To demonstrate the capability of this preparation for chronic, high-resolution physiological monitoring, we performed longitudinal voltage recordings of a subtype of Mushroom Body Output Neurons (MBON05) over a 10-day period. We used the MBON05-specific split-GAL4 line *SS01188* to express the voltage indicator pAce into the pair of MBON05 neurons. Through the optical window, we confirmed that the targeted neuron’s morphology matched the anatomical structure of MBON05 in the FlyWire database (Figure 6A). Neuron morphology remained consistent, with no observable degradation over the 10-day period (Figure 6B), indicating stable optical access over time. Voltage recordings from each session exhibited high signal-to-noise ratios and clearly resolved action potentials with comparable spiking rates across days (Figure 6C). Spike-triggered averages produced nearly identical waveforms across sessions, with no detectable changes in action potential kinetics over the 10-day period (Figure 6D). Together, these results demonstrate stable longitudinal voltage imaging and preserved neural physiology at single-spike resolution over extended timescales.

**Figure 6.**
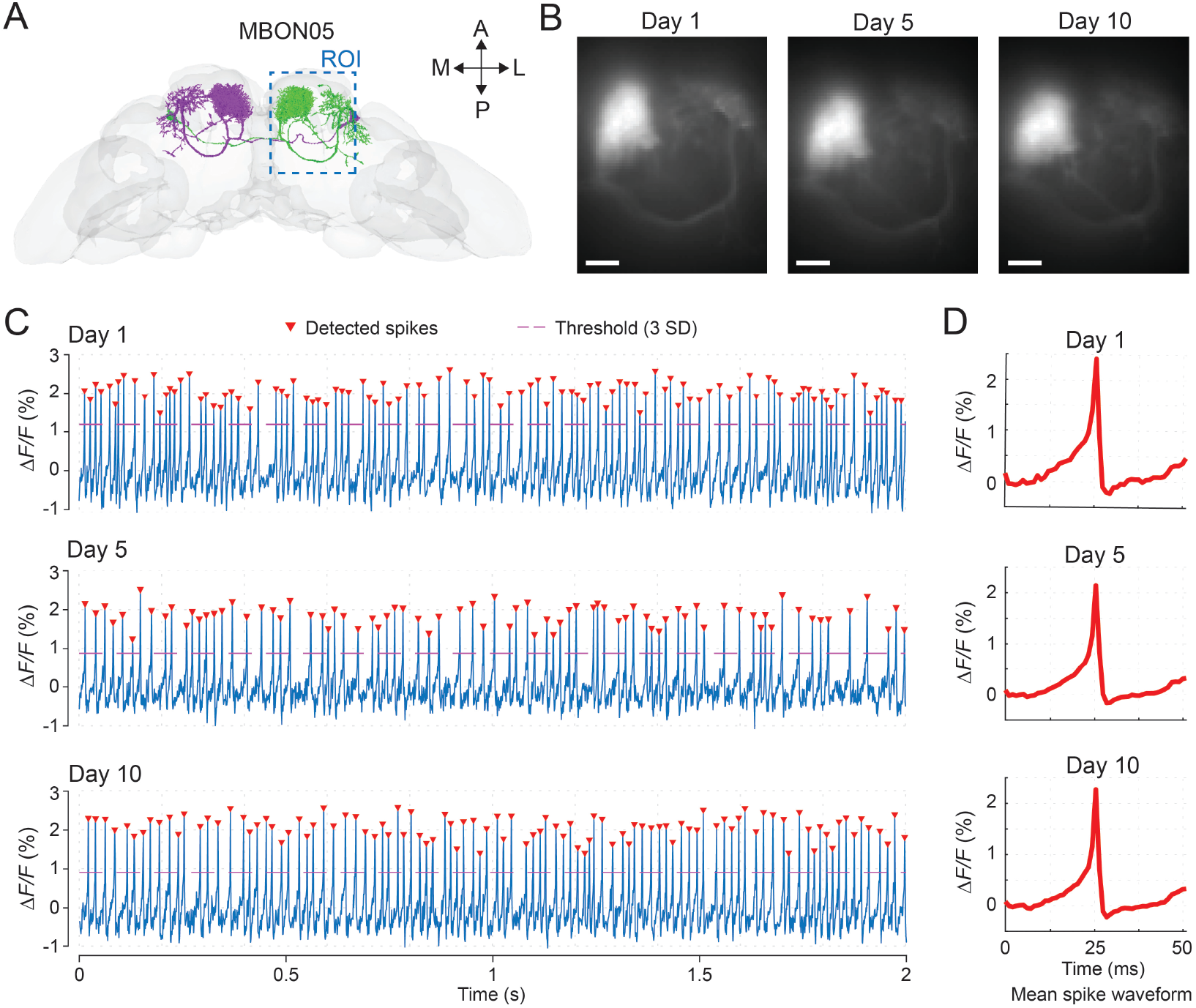
Longitudinal voltage imaging across 10 days. (A) Illustration of the MBON05 neuron morphology from the FlyWire connectome database. Blue dashed line outlines the imaging region-of-interest that covers the dendrite and axon regions of the MBON05 neuron. (B) Fluorescence images show the neuronal structure from the ROI at 1st, 5th, and 10th days post-surgery. Scale bars: 20 µm. (C,D) Example voltage imaging traces (C) and averaged spike waveforms (D) of the neuron at 1st, 5th, and 10th days post-surgery. Red arrows indicate identified spikes that surpassed the detection threshold (3 standard deviations of the whole trace; red dashed lines).

### Two-photon calcium imaging of Mushroom Body Kenyon cells over one week

To demonstrate the reliability of the preparation for population-level functional imaging, we performed longitudinal recordings of odor coding in Kenyon cells (KCs) across 7 days post-surgery. Using the pan-KC driver MB010B-GAL4, we expressed the calcium indicator GCaMP8s in the entire KC population. During imaging sessions, the fly was positioned on a freely rotating, air-suspended ball and odors were delivered through an odor nozzle (Figure 7A). Using two-photon calcium imaging, we successfully tracked the activity of approximately 60 individual neurons per fly across imaging sessions on days 1, 3, and 7 (Figure 7B). Our data indicate that the majority of KCs exhibited consistent odor-evoked calcium transients and stable odor tuning over the week-long period, as shown in three representative neurons (Figure 7C). These results confirm that the surgical and imaging protocols support stable, single-cell resolution tracking of population dynamics over extended timescales.

**Figure 7.**
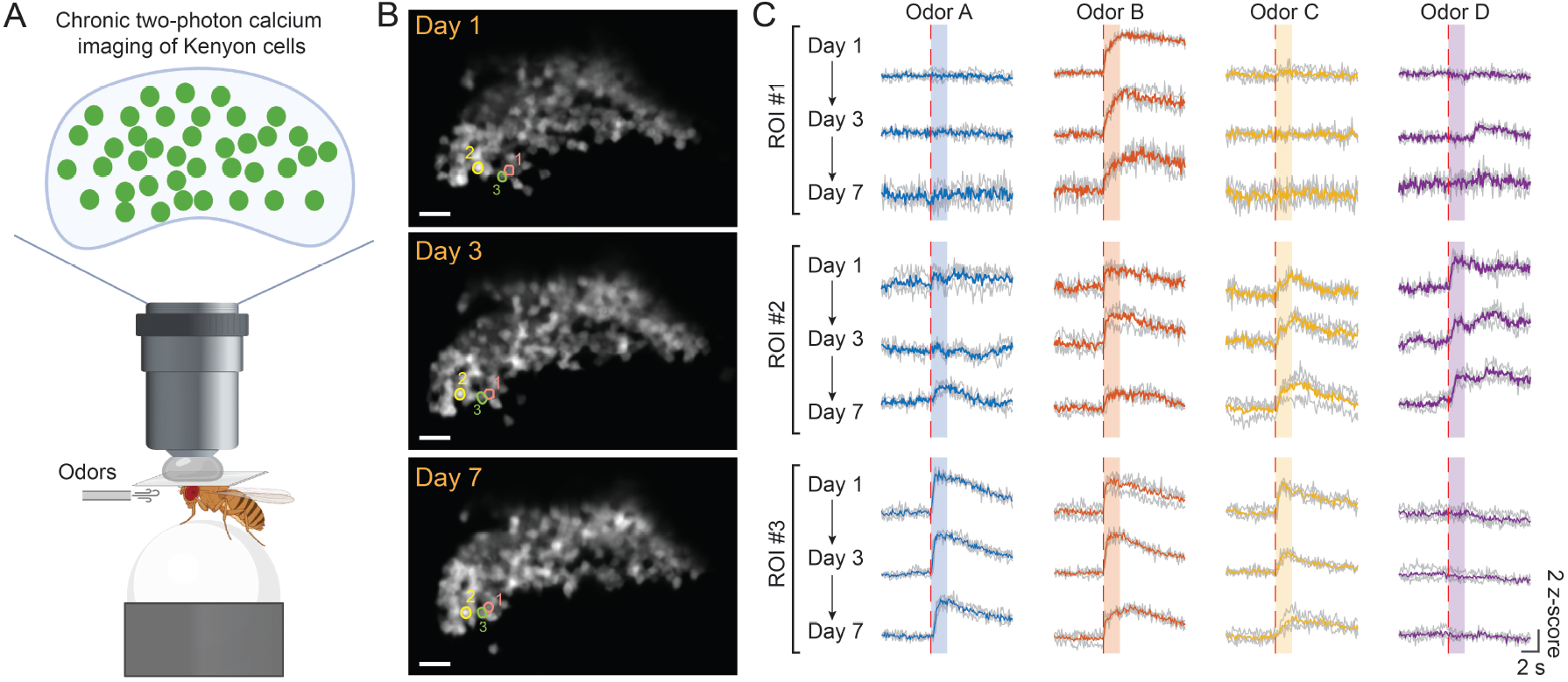
Week-long, two-photon calcium imaging of population activity. (A) Experimental setup for two-photon imaging of odor-evoked activity in the Kenyon cell population. (B) Averaged fluorescence images of the GCaMP expressing KCs at 1st, 3rd, and 7th days post-surgery. Colored circles indicate the three example KCs. Scale bars: 20 µm. (C) Odor-evoked calcium activities from the three example KCs at 1st, 3rd, and 7th days. Dashed red lines indicate the onsets of odor delivery, and the shadow areas indicate the 2 s duration of odors. Solid traces: averaged response from three trials (gray traces).

The raw and analyzed datasets are available at: https://zenodo.org/records/18675613.

## Limitations

Although this protocol enables reliable chronic in vivo recordings, it has several limitations. First, brain motion can occur intermittently. In most cases, the displacement is confined to the xy-plane and can be corrected during post-processing. However, z-axis motion is more challenging to compensate for and often necessitates excluding the affected data. Second, the procedure remains invasive and sensitive to surgical technique, and suboptimal surgery can compromise animal health. For example, low ambient humidity can rapidly desiccate the exposed tissue, adhesive can inadvertently seep into the brain, or tissue damage can occur during dissection or window placement. Neural recordings that exhibit atypical activity patterns (e.g., bursting or abnormally regular spiking) or fail to respond to sensory stimulation should be excluded from subsequent analyses. Third, the window size and location may restrict access to certain brain regions, and the thickness of the glue and coverslip may limit the performance of high-NA objectives or reduce the working distance. Fourth, because the preparation is performed manually, post-surgical survival varies substantially across experimenters. However, with sufficient practice, most experimenters can achieve a survival rate of over 80% for at least a week. Finally, the protocol is optimized primarily for female flies, as males are smaller and, as previously reported^5^, have reduced survival following surgery.

## Troubleshooting

### Problem 1

Ambient humidity levels below 35% or above 60% can interfere with fly mounting and reduce preparation quality. Low humidity increases static electricity, making it difficult to maintain the fly in a natural upright posture on the cooled surface, and increases the risk of tissue desiccation during surgery. High humidity can cause condensation or water droplets to form on the cooling block/washer, which can mix with the epoxy and weaken adhesion.

### Potential solution

- Use a desktop humidifier when humidity is low. Reduce humidifier output or slightly raise the cooling block temperature if condensation forms.
- Maintain relative humidity in the moderate range (typically 35–50%) to prevent desiccation, but low enough to avoid visible condensation.
- Before applying epoxy, use a Kimwipe to remove any water droplets from the washer and cooling surface.
- Minimize direct airflow from vents around the tethering station to stabilize humidity.

### Problem 2

The fly dies immediately after UV curing due to excessive heating, especially when using a different UV light source.

### Potential solution

- Adjust UV exposure time or increase the distance between the UV wand and the fly to prevent overheating.
- Use the minimal amount of glue required for bonding.
- Standardize wand distance and verify curing settings by hardening a test droplet on glass before mounting flies.

### Problem 3

The fly detaches from the pin, and/or the head detaches from the thorax, during surgery or imaging.

### Potential solution

- Ensure the thorax–pin bond has sufficient contact area and that the epoxy is fully cured before proceeding.
- Remove any moisture or condensation from the bonding site (see Problem 1).
- To strengthen the head-thorax fixation, after stabilizing the head, then invert the fly (dorsal side facing the aluminum block), apply approximately 0.5 µL of UV epoxy to each side of the head, and cure the glue.
- If detachment happens frequently, slightly increase curing time (without overheating), clean or replace the pin, and confirm that the epoxy is not partially polymerized.

### Problem 4

Glue seeps into the brain during cuticle resealing, causing acute loss of optical clarity and/or abnormal neural activity.

### Potential solution

- Reduce the volume of sealing epoxy (NOA 68) and apply it along the cuticle rim rather than directly over exposed tissue.
- Avoid pressing the coverslip into uncured epoxy; cure the first layer of glue and then coat a second layer on top before gently placing the coverslip.

### Problem 5

Severe z-motion occurs across frames during imaging, making the data difficult or impossible to correct.

### Potential solution

- The neck is the primary pivot point for head movement. Apply a small amount of additional UV epoxy to the dorsal neck region to rigidly couple the head to the thorax.
- Ensure the acrylic fly holder is securely clamped to the imaging stage and that the stage itself is free of vibration.

## Supporting information

Methods video S1. Video illustration of the fly tethering procedure

Methods video S2. Demonstration of the saline-free surgery procedure (played at 2x speed)

Methods video S3. Release of the tethered fly from the holder

## Resource availability

### Lead contact

Further information and requests for resources and reagents should be directed to and will be fulfilled by the lead contact, Cheng Huang.

### Technical contact

Technical questions about executing this protocol should be directed to and answered by the technical contact, Ruibao Zhu.

### Materials availability

More detailed designs of the mounting platform and *Drosophila* lines are available upon request from the lead contact.

### Data and code availability

The imaging analysis datasets and codes are available upon request from the lead contact.

## Acknowledgments

We thank the members of the Shaw and Taghert labs at Washington University for their helpful discussion. Figures were partially created using BioRender. This work was supported by a Whitehall Research grant (#2024-12-093) to C.H. This research made use of the Neurotech Hub at Washington University in St. Louis.

## Author contributions

C.H. conceptualized the study. R.Z. formulated the method and refined the steps. S.K. designed and built the mounting station. J.Z. and Z.F. collected and analyzed the experimental data. R.Z., S.K., and C.H. wrote the manuscript. All authors reviewed, revised, and approved the final manuscript.

## Videos

**Methods video S1. Video illustration of the fly tethering procedure**.

**Methods video S2. Demonstration of the saline-free surgery procedure (played at 2x speed)**

**Methods video S3. Release of the tethered fly from the holder**.

